# kmerDB: A Database Encompassing the Set of Genomic and Proteomic Sequence Information for Each Species

**DOI:** 10.1101/2023.11.13.566926

**Authors:** Ioannis Mouratidis, Fotis A. Baltoumas, Nikol Chantzi, Candace S.Y. Chan, Austin Montgomery, Maxwell A. Konnaris, George C. Georgakopoulos, Anshu Das, Dionysios Chartoumpekis, Jasna Kovac, Georgios A. Pavlopoulos, Ilias Georgakopoulos-Soares

**Author notes:** Equally contributing authors (Ioannis Mouratidis, Fotis A. Baltoumas, Nikol Chantzi).

## Abstract

The rapid decline in sequencing cost has enabled the generation of reference genomes and proteomes for a growing number of organisms. However, at the present time, there is no established repository that provides information about organism-specific genomic and proteomic sequences of certain lengths, also known as kmers, that are either present or absent in each genome or proteome. In this article, we present kmerDB, a database accessible through an interactive web interface that provides kmer based information from genomic and proteomic sequences in a systematic way. kmerDB currently contains 202,340,859,107 base pairs and 19,304,903,356 amino acids, spanning 45,785 and 22,386 reference genomes and proteomes, respectively, as well as 14,658,776 and 149,264,442 genomic and proteomic species-specific sequences, termed quasi-primes. Additionally, we provide access to 5,186,757 nucleic and 214,904,089 peptide sequences that are absent from every genome and proteome, termed primes. kmerDB features a user-friendly interface offering various search options and filters for easy parsing and searching. The service is available at: www.kmerdb.com.

## INTRODUCTION

Rapid advances in high-throughput technologies, such as next-generation sequencing and massspectrometry, combined with the improvements of modern computer engineering and software development, has facilitated the generation of accurate large-scale reference genomes and proteomes across all taxonomic domains of life (O’Leary et al. 2016; Schoch et al. 2020; Benson et al. 2013). This amount of data has enabled massively parallel comparisons across genomes and proteomes in an effort to annotate them, define coding regions, discover genes and their functions, and reveal insights from genomic regions that have traditionally been considered functionally irrelevant.

Genomes and proteomes consist of sequences of oligonucleotides and oligopeptides, respectively, which can be partitioned into substrings of a fixed length k, known as kmers. Among these kmers, some are conspicuously absent from a given genome or proteome, and are termed nullomers or nullpeptides (Acquisti et al. 2007; Herold, Kurtz, and Giegerich 2008; Koulouras and Frith 2021; Georgakopoulos-Soares, Yizhar-Barnea, et al. 2021) These kmer sequences have been used for various applications, such as quality control, metagenomics classification, phylogenetic analysis, binning, evolution studies, and identification of pathogens (Jagadeesan et al. 2019; Deurenberg et al. 2017; Brandies et al. 2019; Maljkovic Berry et al. 2020). Remarkably, the resurfacing of nullomers in the human genome has been leveraged to detect cancer (Georgakopoulos-Soares, Barnea, et al. 2021; Montgomery et al. 2023), demonstrating their potential for cancer and disease diagnostics. Similarly, quasi-prime kmers have been defined as a set of sequences that are exclusive to a single species and absent from every other known species with an available reference genome or proteome (Mouratidis et al. 2023).

The first attempt to report such patterns was presented by Koulouras et al. with the creation of a database nullomers.org (Koulouras and Frith 2021). However, several limitations exist. The database only covers a limited number of peptide and nucleic minimal absent words, which are a subset of nullomers and nullpeptides. Furthermore, only two reference proteomes and approximately 1,500 reference genomes were considered, limiting its coverage, scope, and applicability. To date, there is no publicly accessible database that offers the comprehensive collection of peptide and nucleic kmers present or absent in each sequenced species’ genome or proteome in an user-friendly and queryable format. Similarly, there is no database providing kmers that are exclusive to each species (i.e., quasi-primes) or kmers absent across all species (i.e., primes), which could have versatile applications. Therefore, a repository where kmer, nullomer, nullpeptide, quasi-prime, and prime sequences can be queried at a large scale has become highly desirable.

In this article, we introduce *kmerDB*, a web-based database built to systematically catalog sets of DNA kmers, nullomers, nullpeptides, quasi-prime, and prime sequences for 45,785 species and 22,386 proteomes spanning all domains of life. The database provides various filter and search options organized in dynamic tables that can be queried and sorted for analysis. Users can investigate kmer patterns across a wide range of genomes and proteomes, examine kmer composition of various lengths for each organism across different taxonomic levels. Reference genomes and proteomes are linked to established publicly available databases such as the ENA Browser (Leinonen et al. 2011), the NCBI Genome Browser (Pruitt et al. 2009), the UniProtKB Proteome database (UniProt Consortium 2023), and InterPro protein families and domains database (Blum et al. 2021).

## RESULTS

### Overall database statistics

Our aim in developing kmerDB is to create a comprehensive repository of genomic and proteomic kmer information that can uniquely characterize each species. We have extracted the kmer and nullomer sequences of each genome and proteome and provide the shortest species-specific oligopeptides and oligonucleotides, termed quasi-primes, as previously outlined by Mouratidis et al. (Mouratidis et al. 2023). The current version of kmerDB comprises 45,785 reference genomes and 22,386 reference proteomes, which can be queried by NCBI and UniProt IDs or by species name. To construct this database, we parsed 202,340,859,107 nucleotides and 19,304,903,356 amino acids across the reference genome and proteome files. The total number of kmer sequences currently present in the database is 90,511,987,219 for reference genomes and 129,729,971,866 for reference proteomes, whereas the total number of nullomers and nullpeptides is 87,117,352,702 and 84,290,713,235, respectively. The number of nucleic and peptide quasi-primes is 14,658,776 and 149,264,442, respectively, and they can easily be retrieved for each species.

The kmer space expands exponentially with increasing kmer length, and for large values of k, the nullomer space encompasses the majority of the possible kmers. This phenomenon is especially pronounced in viruses, which have relatively small genomes and lack many kmers of length greater than seven base pairs (bps) most likely due to their smaller size. Therefore, we only include kmers and nullomers of length up to seven bps for viral genomes in our database. For eukaryotes, archaea, and bacteria, we extract kmers and nullomers of length six to twelve bps. For all available proteomes, we extract kmers, nullomers, quasi-primes, and primes for lengths three to seven amino acids.

We have previously investigated the existence of quasi-primes, oligonucleotide sequences that are exclusive to a single species with a reference genome and absent from all others (Mouratidis, Konnaris, Chantzi, and Chan et al. 2023). We performed a comprehensive search ranging kmer lengths up to sixteen bps and found the first set of quasi-prime sequences at sixteen base pairs, which are provided in the database. Additionally, we have previously examined the occurrence of peptide quasi-primes present in each reference proteome across all species (Mouratidis et al. 2023). No peptide quasi-primes were found for kmer lengths below six amino acids. However, we detected peptide quasi-primes at six and seven amino acids kmer length, which are also accessible in the database. Furthermore, we provide the set of nucleic and peptide primes of lengths of sixteen bps and six and seven amino acids. These are sequences that are absent across all the reference genomes and proteomes, comprising 5,186,757 nucleic primes and 214,904,089 peptide primes.

### The kmerDB interface

A user can navigate the database between Genomes and Proteomes as well as taxonomic groups of bacteria, eukaryotes, archaea, and viruses. The data contained in kmerDB can be accessed through the *Browse* menu located on the kmerDB navigation bar, which enables the user to choose among four domains of life: archaea, bacteria, eukaryotes, and viruses. The user can also specify between genomes and proteomes or use a combination of both criteria. The kmerDB browse page displays a list of genomes and proteomes that can be parsed to match the selected filters (**Figure 2**). The user can further refine the search by opting to select specific species using the NCBI ID, UniProT ID, or the species name, which takes the user to the specific proteome or genome entry page (**Figure 3**). On this page, the user can view the kmers and nullomers/nullpeptides for each kmer length associated with the selected proteome or genome. Registered users can also access quasi-primes for each species. There is also a click option that links to a webpage containing the nucleic and peptide primes. The user can query the kmers or filter by kmer length for individual species (**Figure 4**). For peptide kmers and nullpeptides, kmerDB provides biochemical properties including polarity and charge are calculated and provided. The user can rank and sort kmers based on these biochemical properties. Similarly, for nucleic kmers, kmerDB calculates and provides the nucleotide composition.

**Figure 1:**
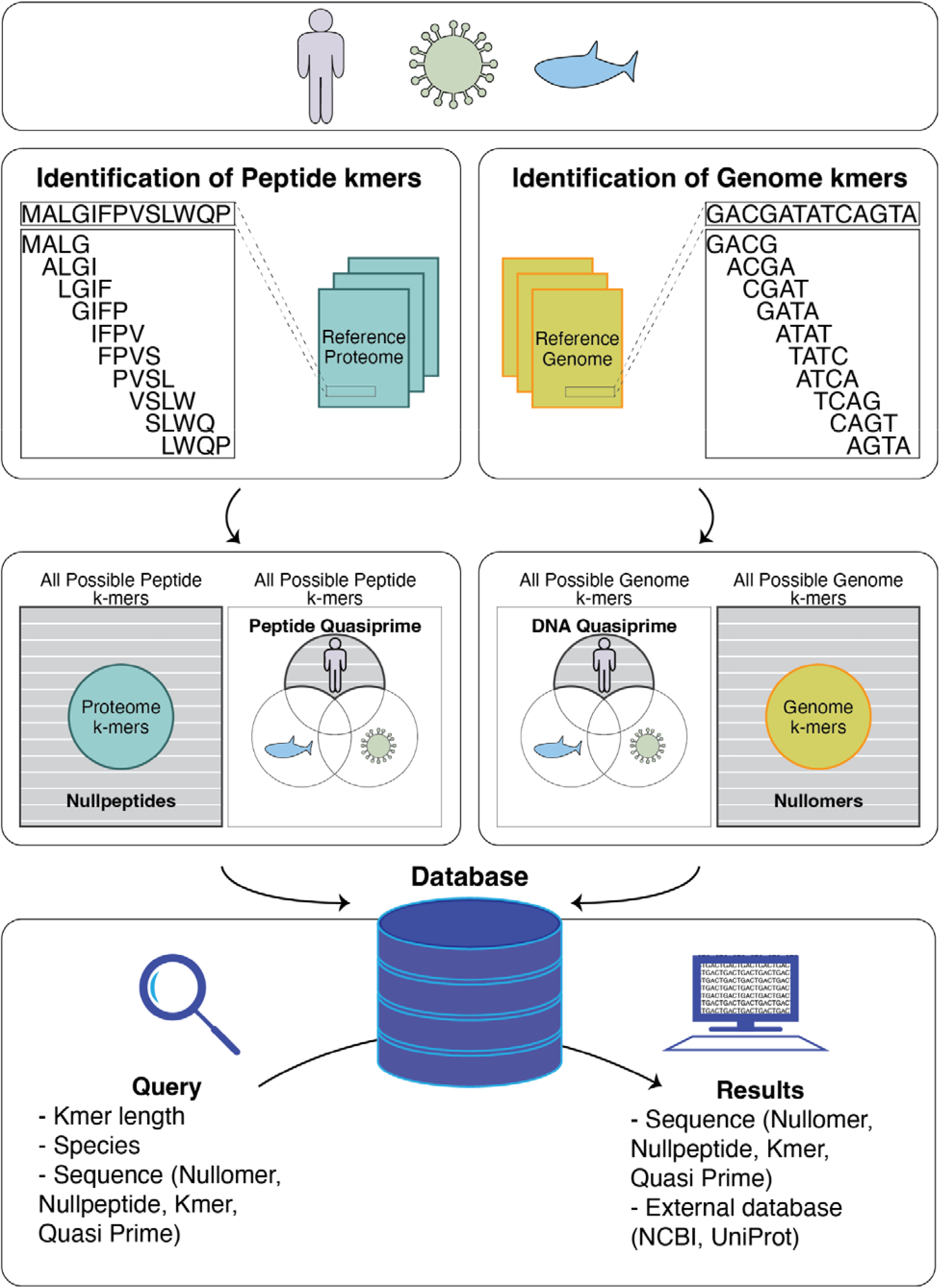
Illustration of the derivation of kmers, nullomers, and nucleic quasi-primes in reference genomes and of kmer peptides, nullpeptides and quasi-prime peptides in reference proteomes. The first step of the process involves cataloging every genome or peptide kmer for each species. The second step involves the derivation of nullomers or nullpeptides. Finally, the set of kmer sequences that are unique to each species are identified. The database encompasses this information for every species and is easily retrievable.

**Figure 2:**
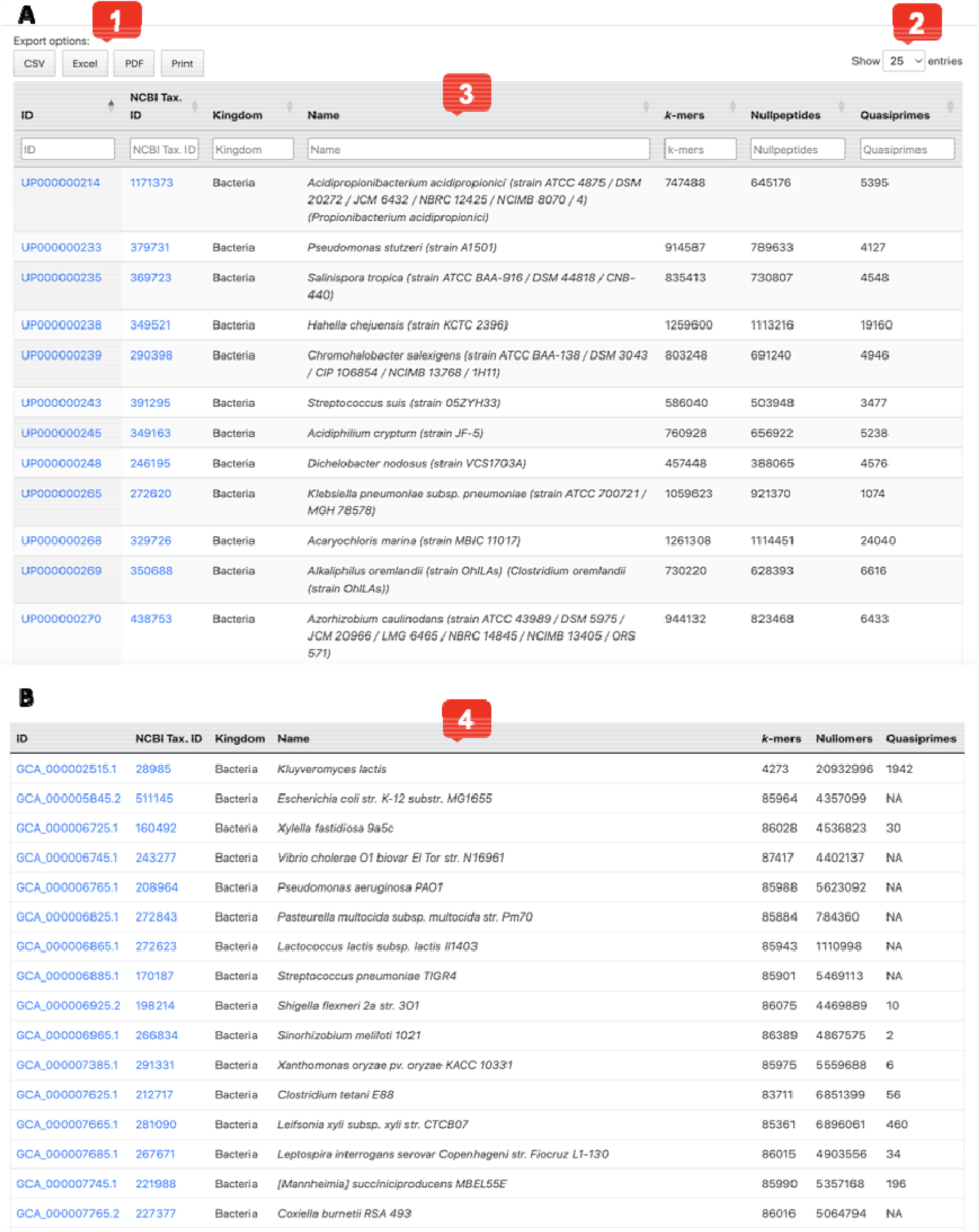
KmerDB Browse pages for genomes and proteomes. **A**. A Table of Entries. Options to download in different file formats (1). By default 25 entries/page is shown (2) with the following columns: UniProt ID, NCBI Taxonomy, Kingdom, species and strain, number of kmers identified, number of nullpeptides detected, and quasi-prime peptides in that proteome (3). **B**. A Table of Entries (by default 20 entries/page) is shown with the following columns: NCBI ID, NCBI Taxonomy ID, Kingdom, species and strain, number of kmers identified, number of nullomers detected, and quasi-prime peptides in that genome (4).

**Figure 3:**
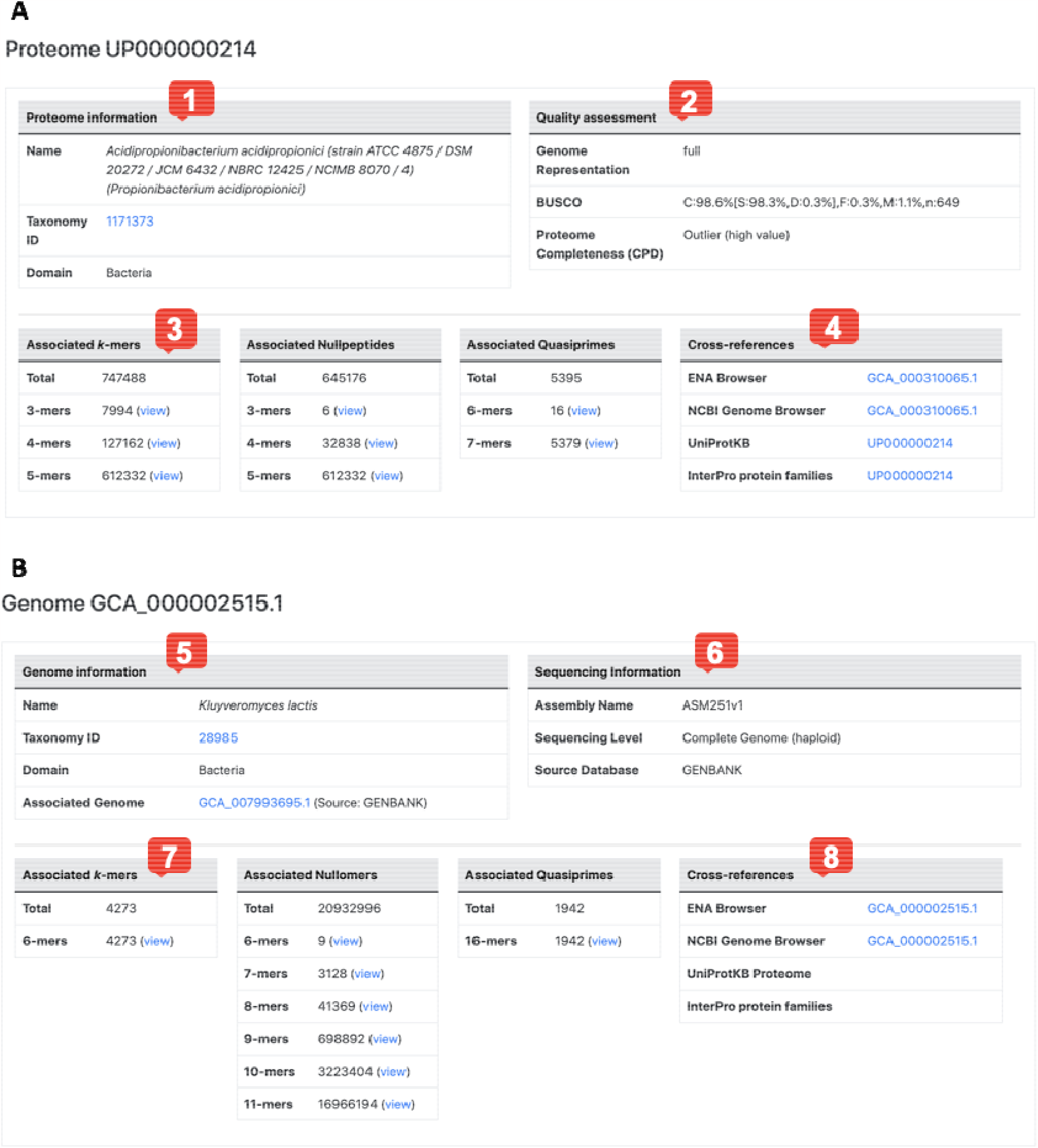
Proteome and genome entry pages. **A**. For each UniProt ID there is an entry page that provides Proteome Information (1) including species name, taxonomy ID, the domain it belongs to, the genome representation, the BUSCO quality metric and its completeness state (3). It also provides clickable options for each kmer length, for proteomic kmers and nullpeptides (3). The user can derive or search for kmers of interest. The entry page also provides cross-references to the ENA Browser, the NCBI Genome Browser, UniProtKB and InterPro protein families databases (4). **B**. For each NCBI ID there is an entry page that provides species name, taxonomy ID, the domain the species belongs to, the associate genome ID, the assembly name (5), the genome completeness state and the source database that was used (6). It also provides clickable options for each kmer length, for genomic kmers and nullomers. The user can derive or search for kmers of interest (7). The entry page also provides cross-references to the ENA Browser, the NCBI Genome Browser, UniProtKB and InterPro protein families databases (8).

**Figure 4:**
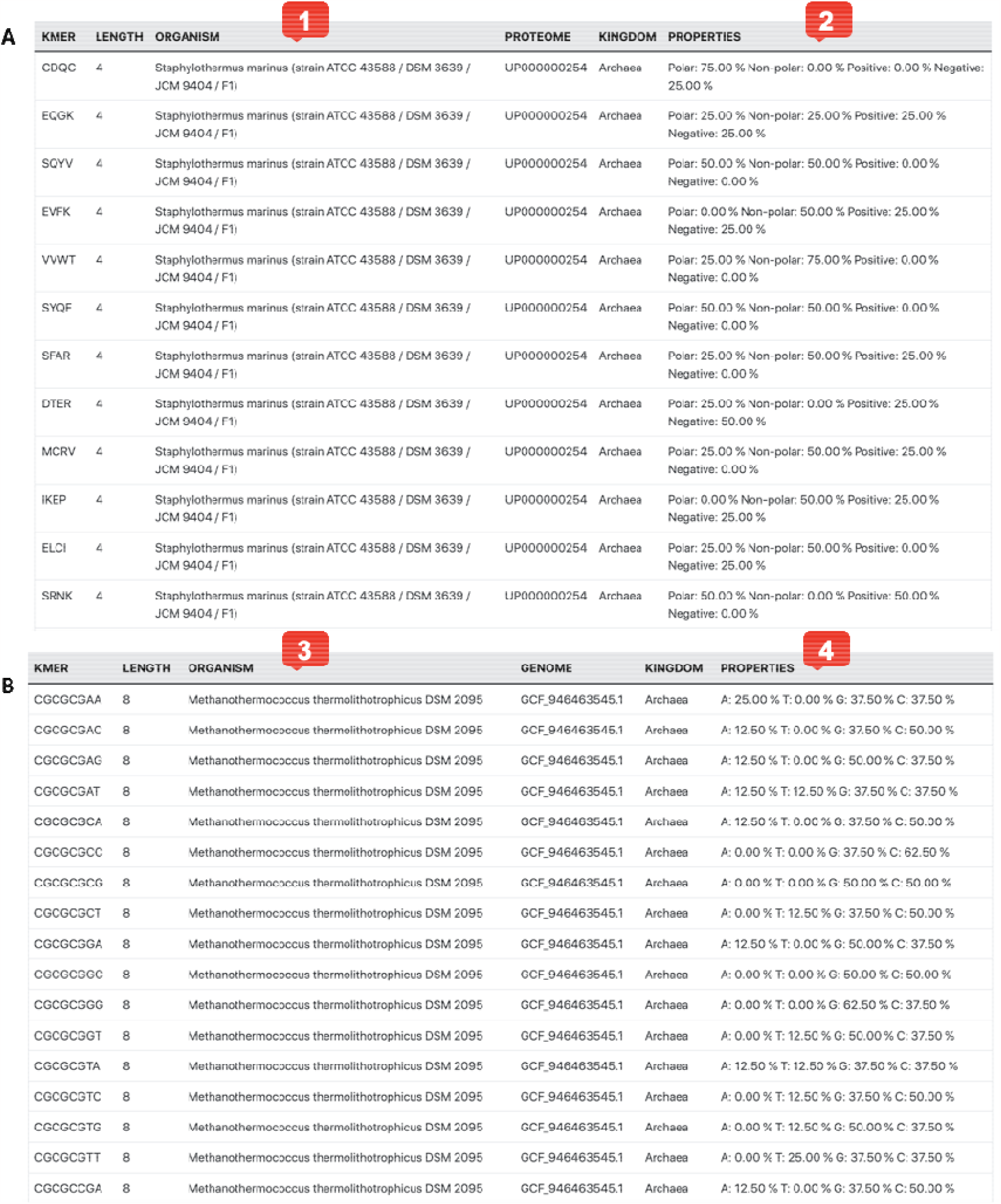
Kmer search page in individual genomes and proteomes for kmers, nullomers, nullpeptides and quasi-primes. **A**. Example search for kmer length of four amino acids in Staphylothermus marinus (1). Information about the peptide content of each kmer is provided under properties (2). **B**. Example search for nullomer length of eight base-pairs in *Methannothermococcus thermolithotrophicus (3)*. Information about the nucleotide content of each kmer is provided under properties (4).

The database is also searchable with a Quick Search Option or an Advanced Search Option. Using the Quick Search Option, the user can search for kmers of interest by UniProt ID or NCBI ID. By using the Advanced Search Option, the user can search by proteome or genome ID, taxonomy ID, organism name, kingdom, number of kmers in a species’ genome or proteome, number of nullomers in a species’ genome, number of nullpeptides in a species’ proteome or number of quasi-primes found in a species’ genome or proteome.

Finally, kmerDB provides links to external genomic and proteomic databases such as the ENA Browser (Leinonen et al. 2011), the NCBI Genome Browser (Pruitt et al. 2009), the UniProtKB Proteome database (UniProt Consortium 2023), and the InterPro protein families and domains database (Blum et al. 2021).

## MATERIALS AND METHODS

### Data retrieval and parsing

Reference proteomes were downloaded from UniProt: (Release 2022_03, 19-Sep-2022). These included reference proteomes for eukaryotes, bacteria, archaea, and viruses (**Supplementary Table 1**). When parsing the proteomes, only the twenty standard amino acids were used, throughout the analyses. Kmer lengths up to and including seven amino acids were studied.

Reference genomes were downloaded from the GenBank and RefSeq databases (Benson et al. 2013; O’Leary et al. 2016) as well as 104 reference genomes from the UCSC genome browser website (Nassar et al. 2023) (**Supplementary Table 1**). Kmer lengths up to and including twelve bps were analyzed to derive kmers and nullomers, whereas sixteen bps was the kmer length at which nucleic quasi-primes were analyzed.

### Definitions

#### Genomic definitions

Let us define the alphabet *L = {A, T, C, G}* representing Adenine, Thymine, Cytosine and Guanine respectively.

We define a ***sequence*** *S= a*_1_*a*_2_*a*_3_.. . *a*_n_ where *a*_*i*_ ∈ *L* for each 1 ≤*i* ≤ *n*.

A ***genome*** consists of a set of sequences over alphabet *L*. In this paper, we use the term kmer to refer to a short sequence *s = b*_1_*b*_2_*b*_3_. .. *b*_k_ of length *k*. We say that a *kmer s* is present in genome *G = { S*_1_, *S*_2_, *S*_3_, . ..,*S*_*L*_ } if and only if there exists *S*_*i*_ *∈ G* such as s is a subsequence of *S*_*i*_ If a kmer s is present in genome *G*, we write *s* ∈ *G*.

A ***nullomer*** of genome *G* is defined as a kmer *s*^*′*^ that is not present in genome *G*, meaning ∄*S*_*i*_ *∈ G* such as *s*^*′*^ is a subsequence of *S*_*i*_ . Therefore a nullomer for genome G is any kmer that is not present in that genome.

For the purposes of this analysis we have considered *X* reference genomes found in the GenBank and Refseq databases. Let *P = {G*_1_, *G*_2_, *G*_3_, …, *G*_x_} the set of all genomes considered. We define a sequence q as a ***quasi-prime*** if and only if there exists *1 ≤i ≤ x* such that *s ∈ G*_*i*_ and *s ∉ G*_j_, ∀*j ∆ i*. Therefore, quasi-primes represent all kmers that are present in only a single genome and absent from every other genome in our database.

A kmer *p* is defined as a ***prime*** in our dataset if and only if ∄ *i* such that *p ∈ G*_*i*_ . Therefore primes represent all theoretically possible kmers that are absent from every genome in our database.

#### Proteomic definitions

Similarly, we define alphabet *L*_*p*_ *= {G, A, L, M, F,W, K, Q, E, S, P, V, I, C, Y, H, R, N, D, T}* representing the common amino acids.

A proteome consists of a set of sequences over alphabet *L*_*p*,_. Proteomic kmers, nullomers, quasi-primes and primes are defined equivalently to their genomic counterparts.

### Nucleic and peptide kmer and nullpeptide detection

Identification of kmers was performed as previously defined in (Georgakopoulos-Soares, Yizhar-Barnea, et al. 2021). For nucleic kmers and nullomers, the lengths of six to twelve bps were used for eukaryotes, bacteria, and archaea, whereas for viruses due to their smaller genome size, the lengths of three to six bps were used. For peptide kmers, oligopeptide lengths of up to seven amino acids were used across all the species. Nullomer and nullpeptide detection was performed as previously described in (Georgakopoulos-Soares, Yizhar-Barnea, et al. 2021) for each species at each kmer length.

### Identification of nucleic and peptide quasi-primes

DNA quasi-prime identification was performed by identifying kmers that were present in each reference genome and nullomers in every other reference genome. Similarly, peptide quasi-prime identification was performed by identifying kmers that were present in each reference proteome and nullomers in every other reference proteome.

Identification of nucleic quasi-primes was performed for kmer lengths of sixteen bp. These are the shortest kmer lengths at which we observe DNA quasi-primes. Similarly, for peptide kmers, we performed quasi-prime identification for kmer lengths of six and seven amino acids, because these are the shortest peptide lengths at which we observe them.

### Database Implementation

The front end of kmerDB is implemented in HTML, CSS, and JavaScript. The back end is supported by the Apache web server and the Slim Framework v. 4.0, with server-side operations handled by PHP and, when required, Python. Genome and proteome metadata are stored in a MySQL relational database. At the same time, kmers, nullomers, nullpeptides, quasi-primes, and primes are organized in prefix tree (trie) data structures, using the Matching Algorithm with Recursively Implemented StorAge (MARISA) Trie implementation and its Python bindings. The kmerDB website layout is designed using the Bootstrap v. 5 framework, jQuery, and the DataTables library.

kmerDB is publicly available through http://www.kmerdb.com. Access to genome and proteome kmers, nullomers, and nullpeptides is open to all users of kmerDB, while access to quasi-primes and primes is restricted.

## DISCUSSION

Herein, we introduce kmerDB, a novel repository that contains kmer, nullomer, nullpeptide, quasi-prime, and prime sequences for 45,785 species with a reference genome and 22,386 species with a reference proteome. While the identification of kmers and nullomers for a species can be obtained with bioinformatic tools (Marçais and Kingsford 2011), for the first time, all kmers, nullomers, nullpeptides, and quasi-primes for each organism are available in a central location, kmerDB. The database provides a user-friendly interactive platform that allows users to select species by name, ID, kmer sequence, or kmer length and provides links to other reference databases, including NCBI for genomic kmer sequences (Pruitt et al. 2009) and UniProt for peptide kmer sequences (UniProt Consortium 2023). The database will be updated regularly to incorporate new reference genomes and proteomes as they become readily available, reflecting the genomic and proteomic diversity of nature.

Here, we outline a few of the potentially highly useful applications of kmerDB across various fields of research. First, DNA quasi-primes are universal and succinct genomic fingerprints for each organism and could therefore be used as detection platforms for pathogens, overcoming the limitations of traditional methods such as cell culture and colony counting that are slow and not applicable to non-culturable species. Second, quasi-primes can also provide insights into evolution and potentially be used as sites of accelerated evolution and sites associated with the development of species-specific traits (Hubisz and Pollard 2014; Mouratidis, Konnaris, Chantzi, and Chan et al. 2023). Third, the identification of nullomers can be used for studies in evolution as a marker of negative selection (Koulouras and Frith 2021; Georgakopoulos-Soares, Yizhar-Barnea, et al. 2021), for pathogen detection or therapeutics (Silva et al. 2015; Pratas and Silva 2021), as anti-cancer agents (Alileche et al. 2012; Alileche and Hampikian 2017), in cancer detection (Georgakopoulos-Soares, Barnea, et al. 2021; Montgomery et al. 2023), as vaccine adjuvants (Patel et al. 2012) or in forensic applications (Goswami et al. 2013). Fourth, kmer information derived from kmerDB can be used for comparative genomics and evolutionary studies (Perry and Beiko 2010; Sims et al. 2009) and for sequence specification such as the identification of CRISPR target sites that are highly specific (Zhu and Cheng 2022). Fifth, prime sequences can be used as genetic barcodes or targetable landing pads in biotechnological applications. Sixth, antibody specificity remains a significant issue, with antibodies usually having cross-reactivity with different proteins (Egelhofer et al. 2011; Bordeaux et al. 2010). Therefore peptide quasi-primescan be potentially used for the design of antibodies with higher specificity.

In summary, kmerDB is a versatile, fast, and high-quality database designed to enable easy access to genomic and proteomic information across species and taxonomies.

## DATA AVAILABILITY

kmerDB is publicly available as a web service at: https://www.kmerdb.com

## AUTHOR CONTRIBUTIONS

G.A.P. and I.G.S supervised the work. I.M., F.A.B., N.C., C.S.Y.C., G.C.G., and I.G.S. wrote the code and generated the schematics. I.M., F.A.B., M.A.K., G.A.P., and I.G.S. wrote the manuscript with inputs from all authors.

